# Prophages in the infant gut are largely induced, and may be functionally relevant to their hosts

**DOI:** 10.1101/2021.06.25.449885

**Authors:** Tamsin A. Redgwell, Jonathan Thorsen, Marie-Agnes Petit, Ling Deng, Gisle A. Vestergaard, Jakob Russel, Hans Bisgaard, Dennis S. Nielsen, Søren Sørensen, Jakob Stokholm, Shiraz A. Shah

## Abstract

**Background:** Bacteriophages are the most abundant biological entity on the planet, and are key components of any ecosystem they are present in. The gut virome is increasingly being implicated in disease states although these studies largely focus on lytic phages in adults. Here we identify prophages from a large infant cohort and investigate their potential functions.

**Results:** We identified 10645 vOTUs from 662 metagenomes. No core virome was found: the most prevalent vOTU was identified in 70% of the samples. The most abundant and prevalent group of phages are a novel group closely related to *Bacteroides* phage Hanky p00’. Functional annotation of this group revealed the presence of genes in the dDTP-L-rhamnose pathway, possibly involved in the production of capsular polysaccharides. We also found an abundance of diversity generating retroelements in the phages. Additionally, paired virome data allowed us to show that the majority of prophages are induced in at least one sample and that this is not affected by the use of antibiotics in the 4 weeks prior to sampling.

**Conclusions:** Prophages in the infant gut are largely unique to the individual and not shared. Most of them appear to be induced and so may be key drivers in shaping the bacterial microbiome. The most abundant group of phages are novel, and possess elements that may allow them to maintain differentially susceptible subpopulations of their host bacterium; whilst also containing diversity generating retroelements that could expand their host range. Prophages are important components of the infant gut that may have far reaching influences on the composition and function of the microbiome.

## Introduction

The human gut microbiome is a complex and diverse environment that is established soon after birth and becomes increasingly diverse over the following few years, until a stable ‘adult-like’ composition is reached during the preschool years [1–5]. Within this period of maturation, a number of factors have been associated with differential development of the gut microbiome. The most well studied of these is birth mode[6, 7], but factors such as medication use[8], diet [2, 9, 10], rural or urban environments [11, 12], and the influence of siblings and pets have also been found to have an effect on bacterial composition [13]. The early bacterial microbiome has been associated with a range of disease outcomes in later life including asthma [14], allergy [15], and inflammatory bowel disease [16]. Our understanding of the development of the bacterial microbiome and its role in disease states has greatly improved over recent years, and we now recognise it as a key component of these states. Whilst much work has been done on the bacterial component, recent studies have also shown the viral community to be altered in certain diseases [17], suggesting that they may also play a role. The viruses that make up a large proportion of the microbiome are collectively referred to as the ‘virome’, and in recent years their contribution to this environment has been increasingly investigated although comparatively much less is still known.

Though the virome consists of both bacteriophages (phages), as well as eukaryotic viruses, we will refer to phages only when talking about the virome here. Shifts in the phage composition of the gut have been associated with an increasing number of diseases such as Crohn’s disease and Ulcerative Colitis [17] as well as other conditions such as arthritis [18]. These phages are thought to be virulent, but previous work has suggested that the gut virome contains a significant proportion of temperate phages as well [19, 20]. Temperate phages can integrate into the genomes of their bacterial hosts and be maintained through generations either as a part of the host genomes or as a plasmid-like object; but can also return to the lytic cycle if the bacterial host is stressed by external factors [21]. Up to 20% of the phages found by Reyes *et al* in faecal samples from adults were predicted to be temperate [20], and previous studies have found that multiple strains of bacteria from the gut contain prophages, suggesting that lysogeny is a widespread phenomenon [22]. Whilst virulent phages infect and kill their host cell, temperate phages that have become incorporated into the genome of their host may therefore be under a selective pressure to provide something useful to their bacterial hosts.

Temperate phages provide an opportunity to influence the metabolism and function of their bacterial hosts by harbouring morons [23]. These genes are not essential for the phage itself, but may benefit the bacterial host by providing additional functionality [23]. These can include antibiotic resistance genes [24–26], and toxins or virulence genes that increase the fitness of pathogenic bacteria during infection [27–29]. The presence of a prophage within a bacteria also has the added benefit of being protective from infections. Superinfection exclusion is a method by which prophages prevent infection of their host by related or more distant phages [30–32]. This is achieved by a variety of methods including alterations to the cell membrane [33, 34], repressor-based immunity [35], and inhibition of DNA translocation into the cell cytoplasm [36, 37], amongst others. Whilst there are a number of potential benefits to the bacterium from the possession of a prophage, negative effects have also been identified in some cases. For example, in a monoxenic mouse model system the carriage of lambda prophage in *Escherichia coli (E.coli)* was detrimental to the host bacterium due to frequent reactivation of the prophage [38]. Additionally, in *Streptococcus pneumoniae* the carriage and expression of prophage element Spn1 has been shown to be detrimental to the fitness of the pathogen, by reducing its ability to colonise the nasopharynx [39]. Whether their effects are positive or negative, a growing body of evidence points to temperate phages being key components of the gut microbiome that have the ability to greatly influence the host assemblage diversity and functional potential [20, 22, 40, 41].

In this work we utilise a set of 662 infant metagenomes coupled with viromes to identify and characterise the temperate phages in the gut at 1 year of age and evaluate degree of induction. The samples originate from subjects of the Copenhagen Prospective Studies on Asthma in Childhood 2010 (COPSAC_2010_) cohort, an ongoing mother-child cohort study followed since pregnancy and throughout early life with exhaustive phenotyping and sample collection. By using the paired metagenome and virome data we are able to address the role of prophages in this system; we were able to estimate if these phages are actively induced and whether this is affected by external variables such as antibiotic usage, and start to address the question of what their potential function might be through the analysis of accessory genes. Our work supports the current hypothesis that virome diversity is high in early life due to the induction of prophages and goes on to start to address the question of what their role in this environment might be.

## Methods

### Sample Collection

Samples were collected from the COPSAC_2010_ mother-child cohort that forms the basis of an ongoing cohort study of 738 pregnant women and 700 children that have been followed from week 24 of pregnancy. The infant faecal samples studied here were collected at 1 year of age either at the research clinic or at home by the parents following detailed instructions. All samples were mixed with 1mL of 10% glycerol broth at the laboratory and stored at −80°C until use.

### Ethics

The study was approved by the Capital Region of Denmark’s Local Ethics Committee (H-B-2008-093) and the Danish Data Protection Agency (2015-41-3696). The study was conducted following the guiding principles of the Declaration of Helsinki. Before enrolment, parents gave their oral and written informed consent.

### Metagenome and Virome Sequencing and Assembly

The same samples were used for both metagenome and virome sequencing. For virome sequencing they were filtered through a 0.45 μm pore size PES filter (Minisart® High Flow Syringe Filter, Sartorius, Göttingen, Germany). For the metagenome sequencing DNA was extracted using the method of Mortensen *et al* [42] and the DNA was sequenced, and genomes assembled as described in Li *et al* [43]. Sequencing reads have previously been deposited in the SRA under the accession: PRJNA715601. DNA for sequencing of the virome was extracted and sequenced as previously described in Deng *et al* [44]. The virome sequencing reads will be deposited upon submission.

### Putative Prophage Contig Identification, and classification

Putative prophage sequences were identified using a combination of two methods: DeepVirFinder (v1.0) [45] and VIBRANT(v1.0.1) [46]. Assemblies from all 662 metagenomes were concatenated into a single pool and filtered to those above a minimum length of 4 kb. These assemblies were run through DeepVirFinder (v1.0) using default parameters after creating models from all known phage genomes, downloaded from the Millardlab database in September 2019 [47]. Resulting contigs were filtered to include only those with a p-value of <0.05 after FDR correction. The assemblies were also run through VIBRANT (v1.0) using the nucleotide input and parallelisation options. Only those predicted phages of ‘medium quality’ and above were used further. Both sets of output were compared to the Refseq+plasmid database with a cut off of 95% identity using MASH (v2.2) [48] and any contigs with matches that might be contamination were removed. Resulting contigs from both methods were then combined and dereplicated at 95% ANI with dedupe2.sh [49]. This set was also run through CheckV(v0.7.0) [47] to give the details required for the minimum information about uncultivated viral genomes (MIUViG) [50].

Whilst measures have been taken to remove potential bacterial contamination in these sequences by applying appropriate cut-offs with the tools used, there is always the possibility that some bacterial data still remains and this should be considered in any future analysis. Additionally, whilst the phages identified in this work are referred to as prophages, it is important to remember that this assignment is putative, as the discovery of virulent phages is still possible from metagenomic data.

Predicted prophages were clustered using a network analysis performed with vCONTACT2 (v0.9.8) [51], using the RefSeq. 88 database, with all other sequenced bacteriophages included using the Millardlab database as of January 2021[47]. The resulting network was visualised in Graphia (v2.1) [52] with nodes of vOTUs identified in this work coloured in blue and those that were reference sequences coloured in grey, key viral families were manually annotated.

### Prophage Annotation and Functional Analysis

Sequences were annotated using prokka (v1.14.5) [53] and a custom database made from all phage genes using the Millardlab database from September 2019. The –add-genes and –locus-tags options were also used. Resulting amino acid files were clustered at 90% identity using CD-Hit (v4.8.1) [54] and representative sequences from each cluster were analysed using EggNOG-mapper(v2.0) [55] and default parameters. Phage lifestyle prediction was calculated in the same way as Cook *et al* [56] using HMM profiles for proteins that indicate a temperate lifestyle, and hmmscan (v3.3) [57].

### Identification of previously isolated temperate phages

The genome sequences of a set of *E. coli* temperate phages that were isolated from the same infant faecal samples used here [58], were used to evaluate how well the temperate phage identification methods worked, and as reference genomes for comparison. To identify if any of the predicted sequences were those of the previously identified coliphages the reference genomes were dereplicated at 95% ANI to match that of the predicted contigs. The dereplicated contigs and the prophage contigs were then mapped against each other with minimap2 using the -asm20 option [59]. The sequences of any hits with >70% of the target contig covered was then extracted and clustered using Cluster_genomes.pl (v5.1) [60] to agglomerate contigs that were >95% similar over 90% of the genome a cut-off previously accepted as the same genome [61], and the longest representative of the cluster was kept as the representative sequence.

### Distribution of prophages in individuals

A set of reference contigs was constructed using the vOTUs identified from the metagenomes. The genomes of any remaining temperate coliphages isolated from the samples that had not been identified as vOTUs using the method described above were also added. Additionally, a set of 249 reference crAss genomes were downloaded from the dataset constructed by Guerin *et al* [62], to try and capture the diversity of the crAss-like family of viruses that are abundant in the gut. This set of predicted phages, coliphages and crAss phages were then dereplicated at 95% ANI using dedupe2.sh to remove any remaining redundancy [46].

Trimmed and QC’d reads from individual metagenomes were mapped against the set of reference contigs using bbsplit.sh with random mapping of ambiguous reads, and a minimum identity of 0.95, the covstats option was also implemented [49]. A contig was considered to be present in a metagenome if there was coverage of >=1X across >=70% a similar cut off used by Roux *et al* [61]. Abundances were then calculated as counts per million (CPM). To determine how often the phages are present in the metagenomes a binary presence/absence matrix was used so that extreme outliers in abundance would not skew results. The sum of the presence for each phage was calculated and sorted to identify the phages present in the most metagenomes. Alternatively, to identify how many prophages appeared in each child, the sum of presence in each metagenome was calculated.

When characterising the distribution of crAssphages in the samples the reference genomes along with any vOTUs that clustered together with them in vCONTACT were considered crAss-like phages. The sum of presence of this subset of phages was calculated from the presence/absence matrix.

### Host Prediction

Bacterial hosts of the phages were predicted using CrisprOpenDB (v1.0) [63] with 1 mismatch allowed, and the host with the most prophages predicted to infect them were identified in R and the host with more than 1% of the prophages predicted to infect them were visualised.

### Functional Analysis of the most abundant prophage cluster

Proteins from all members of cluster1819 were extracted and clustered at 90% ANI using CD-Hit [52] to remove redundancy. These were then analysed with EggNOG-mapper(v2.0) [55] as previously described to assign Clusters of Orthologous Groups (COG) categories. Additionally, resulting Kegg Orthology (KO) codes were mapped onto metabolic pathways using KEGGmapper [64].

### Phylogenetic Analysis of the most abundant prophage cluster

An initial blast search showed that *Bacteroides* phage Hanky p00’ (Hankyphage) was the only phage with significant sequence similarity to the cluster members: the high quality vOTU_03578 was used as a representative of the cluster and had 99.90% identity and 74% query coverage with *Bacteroides* phage Hanky p00’; therefore, putative terminase genes from all cluster members were identified through analysis of the Hankyphage p00’ genome. The terminase protein sequence of Hankyphage p00’ was downloaded and used to identify the same protein in the cluster members using HMMsearch [57], as no protein had been annotated as such. The protein sequences were aligned with MEGA(v10.1.8) and a maximum likelihood tree was also produced in MEGA using the JTT model and 100 bootstraps [65]. The tree was visualised and manually coloured in iTOL [66].

### Identification of Diversity Generating Retroelements in the most abundant prophage cluster

Diversity generating retroelements were identified in members of cluster1819 by using both the myDGR web server [67] and MetaCSST(v1.0) [68] tools. To predict whether the target genes identified were putative tail fibre genes, as has previously been suggested [69], the proteins from all members were analysed with PhANNs(v1.0.0) [70] and the most significant hit to a tail fibre gene was carried forward. These were then compared with the results from the previous tools.

### Determining ‘active’ prophages

Utilising the paired nature of the metagenome and virome sequencing of the samples allowed for a novel exploration of whether the predicted prophages were induced at the time of sampling. Individual virome sample reads were mapped against the original metagenome assemblies containing the prophages that had been excised by the prediction tool VIBRANT [46]. A total of 4291 prophages were able to be used in this analysis. Using the predicted coordinates of the prophages, the coverage of each background bacterial and predicted prophage region of an assembly was extracted from a bam file that had been sorted and indexed using samtools [71]. For this analysis it was assumed that there was only a single prophage region per contig; there were only a minority of cases where multiple prophage regions had been predicted for a contig and had passed the quality cut-offs used in this work. The assumption of a single prophage region may lead to a small number of false negatives in the induction analysis. A small number of virome samples were also mapped against three large chromosomal contigs that were not predicted to contain any prophages as a negative control. The number of reads mapped to sections of 40kb (the mean size of prophages in this work) were extracted from different regions of the assembly, in the same way as described above to mimic the presence of a prophage and allow us to test for induction in these negative controls.

To determine statistically if prophages were induced the number of reads mapped to the bacterial part of the assembly and the number of reads mapped against the predicted prophage part were tested for a binomial distribution using pbinom in R(version 3.6.1), and the resulting p-values were corrected for multiple testing using Bonferroni method. A linear model was used to test if the fraction of children where a prophage was significantly induced associated with the mean reads per kilobase (RPK) of the prophages to make sure that higher significance was not the purely result of an assembly that had more coverage. The induction significance was compared across predicted hosts of the prophages, and within *Bacteroides* phages those that belong to cluster1819 were compared against the rest. The induction significance of cluster1819 phages was also compared to all other prophages.

## Results

### Identification and Classification of novel Prophages in the Infant Gut

From the 662 infant metagenomes obtained at age 1 year, we identified 10645 vOTUs after dereplication (Supplementary Table 1).

We grouped these viral contigs into approximate genus or subfamily-level classifications using vCONTACT2 and included the genomes of all sequenced bacteriophages as references, resulting in 2934 clusters with 364 singletons and 2221 outliers. Of all the clusters identified 953 were comprised solely of vOTUs identified in this work and of these 953 clusters, 177 were made up of a single member (**Fig 1**). Here reference genomes are in grey and vOTUs identified in this work are in blue to highlight their distribution throughout viral clusters. Key viral families and groups of interest in this paper have been annotated (**Fig 1**). The hosts of 65% of the vOTUs could be predicted; the most common assignment at the genus level was *Bacteroides* (12.2%), followed by *Salmonella* (6.3%) and *Bifidobacterium* (5.8%) (**Fig 2A**) Previous work using these samples were able to isolate 35 *Escherichia* temperate phages, and sequenced their genomes [58]. There were five vOTUs that shared significant similarity with these isolated phages (**Table 1**). The longest representative from each cluster was kept as representative, resulting in four predicted sequences being replaced with the isolated phage genomes and one isolated phage genome replaced by a predicted phage genome. The ability to identify five vOTUs with similarity to previously isolated phages may reflect a level of mosaicism or alternatively, microdiversity between this group of closely related phages. Both of which are known to cause problems with assembly and may explain why more were not found [72, 73]. It may also be a reflection of the fact that the phages able to be isolated may not be abundant enough in the sequence data to assemble fully.

**Figure 1.**
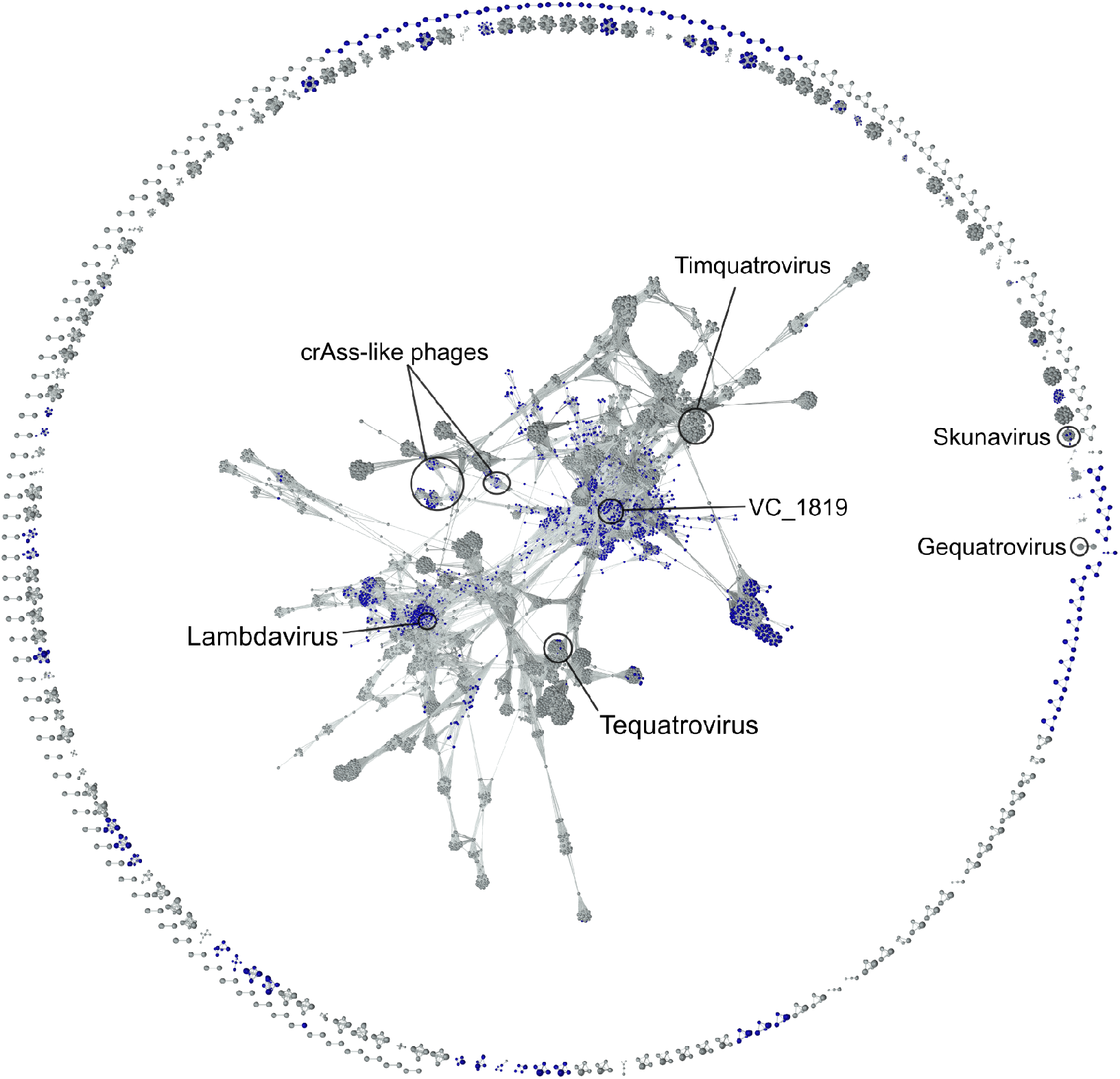
vCONTACT2 network analysis of vOTUs from this study and a database of phage genomes extracted from millardlab.org in January 2021. vOTUs identified here are coloured in blue and reference genomes are grey. The largest and key viral families have been annotated.

**Figure 2.**
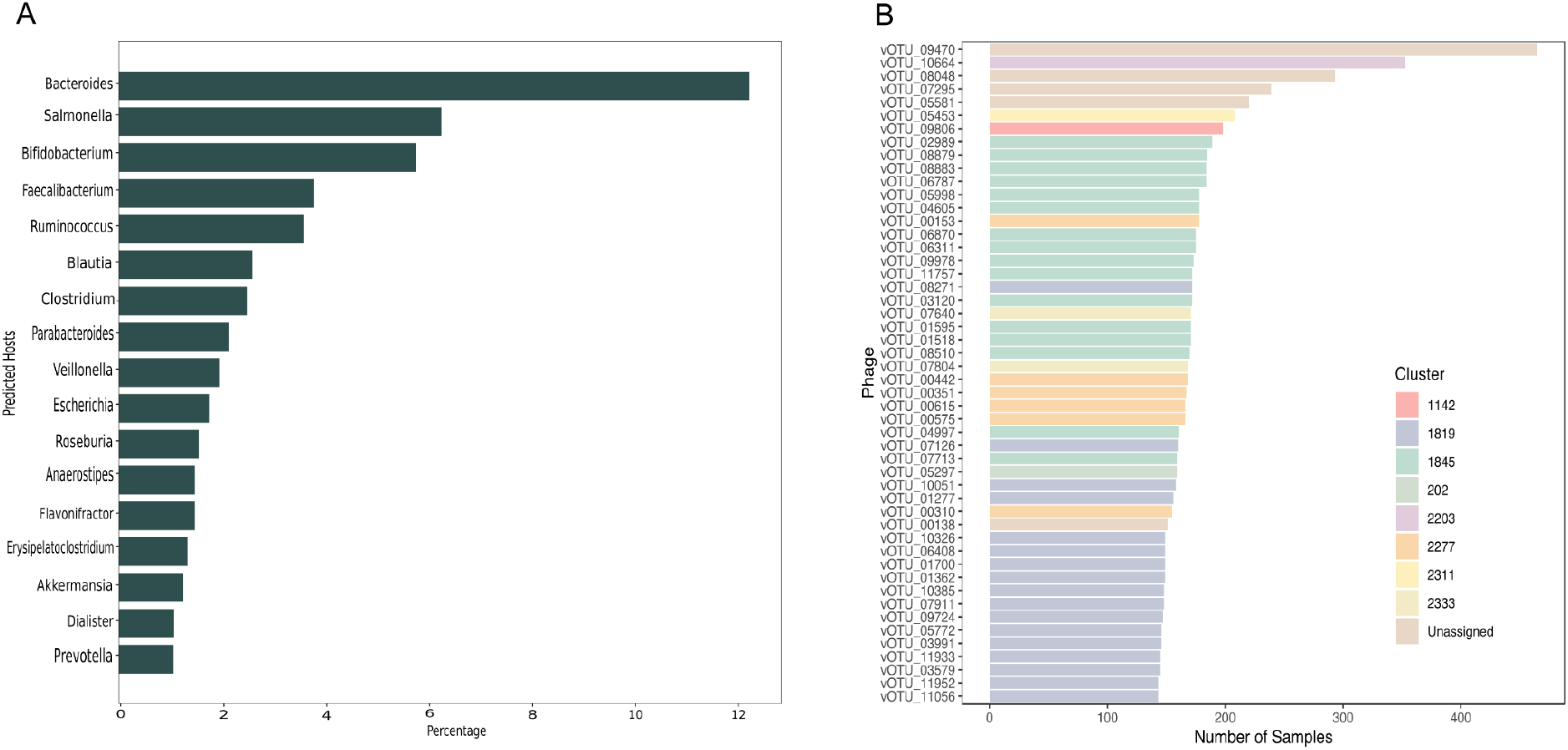
A) Host prediction for the putative prophages. Only those that make up greater than 1% of the predicted hosts are shown. B) Top 50 most prevalent vOTUs coloured by their viral cluster.

**Table 1.**
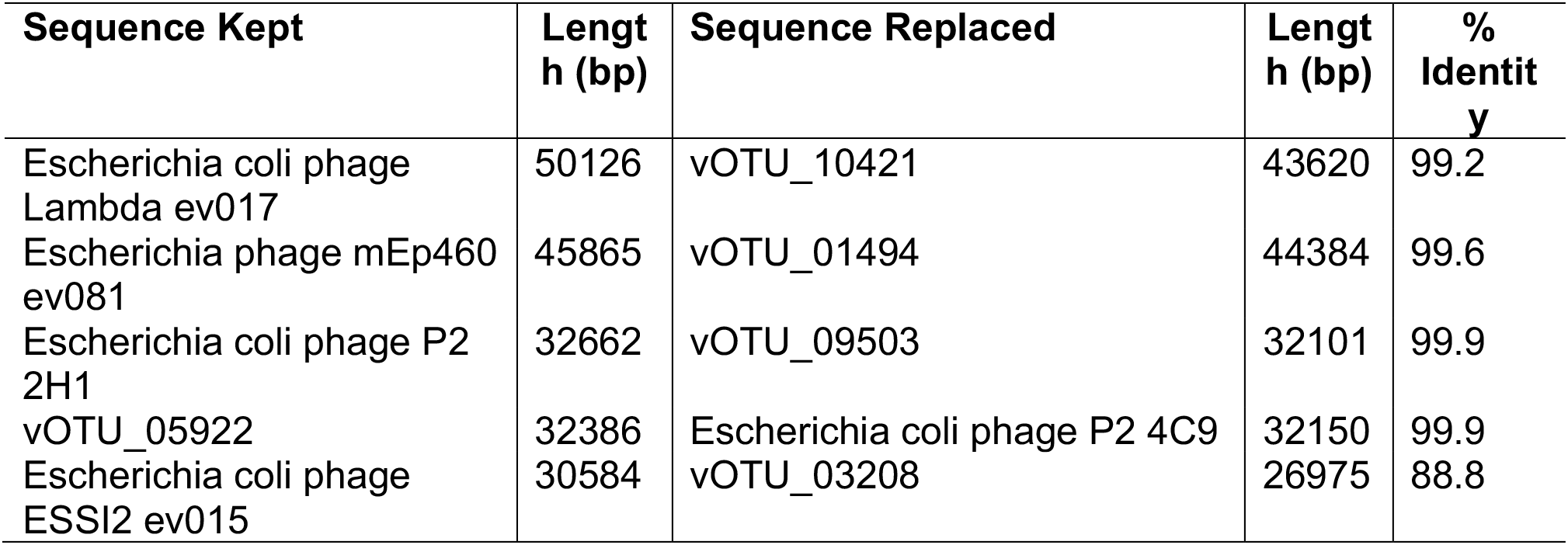
Percent identity of vOTUs identified in this work, and Escherichia coli temperate phages isolated in previous work.

### Abundance Analysis Suggests no ‘Core’ Virome is Established in Infants

The distribution of the number of phages in each sample shows that the average number of prophages per child is ~100, with values as low as four and as high as 400. This is corroborated by the finding that no single phage vOTU was found in all of the children. The most widespread vOTU was found in 71% of the children (**Fig 2B**), far below the 95% cut off used in this work to designate a phage as ‘core’. Using the 50% cut off that has been used in previous work for the same designation [74] results in one additional vOTU. Whilst no individual phage could be found in all samples, the top 50 most widespread phages are spread between only eight viral clusters (excluding those without an assigned cluster) (**Fig 2B**), showing more conservation of the genus/subfamily level than of individual viral contigs.

CrAss-like phages were present in 195 (29%) of the samples sequenced, and if a sample had crAss-like phages identified, it was likely to possess only one type, as only 38% of the crAss-positive subjects contained two or more types. The identification of 109 vOTUs that clustered together with the reference crAssphage genomes has also expanded our knowledge of crAssphage and crAss-like phages.

### The Functional Potential of Viral OTUs showed no significant patterns on the individual phage level

The percentage of all proteins involved in the different COG categories showed that the majority (64.9%) of proteins were assigned to category S – those of Unknown Function. Followed by categories reflecting viral replication – Replication, Recombination, and Repair (12.9%); Transcription (6.6%), and Cell Wall/Membrane Biogenesis (3%). Other categories were present in very small percentages of the total protein amount.

### Cluster1819 is the most abundant phage cluster and contains an abundance of DGRs and morons

Cluster1819, containing 82 members, was found to be the most abundant in the samples (**Fig 3A**) and within the top 50 most widespread vOTUs it was one of the two most prevalent clusters represented and so was characterized in more detail. Phylogenetic analysis of the large terminase gene revealed a single relative: *Bacteroides* Hankyphage p00’ (Accession BK010646) (**Fig 3B**). The bacterial host for Hankyphage was previously identified as a *Bacteroides* which is the same as the predicted host for many members of this cluster [66]. However, there are variations on this with some vOTUs predicted to infect *Prevotella* and *Butyricimonas*.

**Figure 3.**
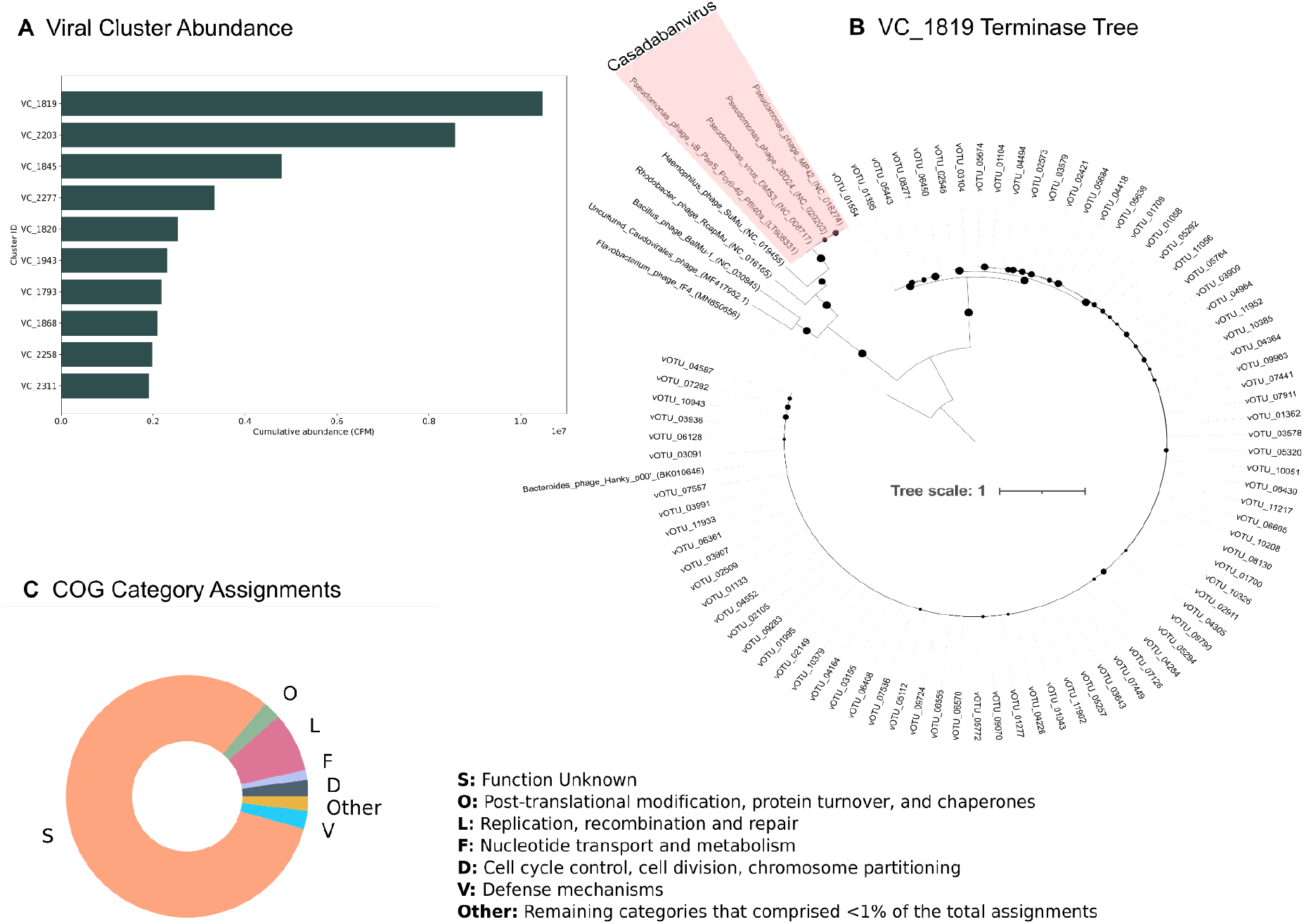
Characterisation of cluster1819. A) Cumulative abundance of the top 10 most abundant viral clusters. B) Phylogenetic tree based on a terminase protein alignment for members of cluster1819 and including the Bacteroides phage Hankyp00’. Only bootstrap values >70% are shown as circles. The next most closely related phages were used as an outgroup due to their limited genome similarity. Any known taxonomy has been highlighted in red. C) COG Category assignments for proteins belonging to members of cluster1819. The ‘other’ group is made up of those categories that comprised less than 1% of the total assignments.

The combination of MetaCSST and MyDGR identified 53 diversity-generating retroelements (DGR) in the cluster, an element that was found to be present in Hankyphage. Of the cluster members 49/82 were found to contain a DGR, as some contained multiple, and the target sequences were used to predict the gene it would generate diversity in. PhaNNs was used to predict the structural genes for the cluster members including the tail fibre genes, commonly a target for DGRs; when the target gene sequences were compared to the structural gene predictions all target genes were predicted to be tail fibres.

The majority of identified proteins in cluster 1819 belong to category S – those of unknown function (**Fig 3C**). This is followed by category L and represents proteins involved in replication, recombination, and repair; categories O and V are equally abundant and represent those proteins involved in post-translational modification, and defence mechanisms respectively. Cell cycle control and nucleotide transport and metabolism are also categories of note. Combining these COG codes with the KEGG pathway database gave a much clearer overview of what pathways could be affected by these phages. These pathways included dTDP-L-rhamnose biosynthesis, and menaquinone biosynthesis amongst others.

### Estimation of the proportion of ‘active’ prophages

Next, we used the results of the virome reads mapped against the large metagenome contigs containing prophages, to discover prophages that were potentially induced and present in the samples as free phages. A subset comprising 4291 of the predicted prophages were able to be tested for induction via a read mapping approach, as they had been excised from a larger assembly in the prediction process, thus allowing for a comparison against the bacterial background. We quantified and tested induction as a degree of preferential mapping of virome reads inside the predicted bounds of the prophage compared with the rest of the contig; for examples see **Fig 4A+B**. The results of this approach showed that induction is a widespread phenomenon; 4041 (94.2%) of the prophages were induced in at least one sample and remained significantly so (p=<0.05) after Bonferroni correction, resulting in 4.59% significant prophage-sample pairs, see **Fig 4C**. Only 250 prophages were never found to be significantly induced in any sample, see **Fig 4D**. When only considering contigs attracting 100 reads or more in a sample, 83418 out of 321232 pairs (26.0%) were found to be significant.

**Figure 4:**
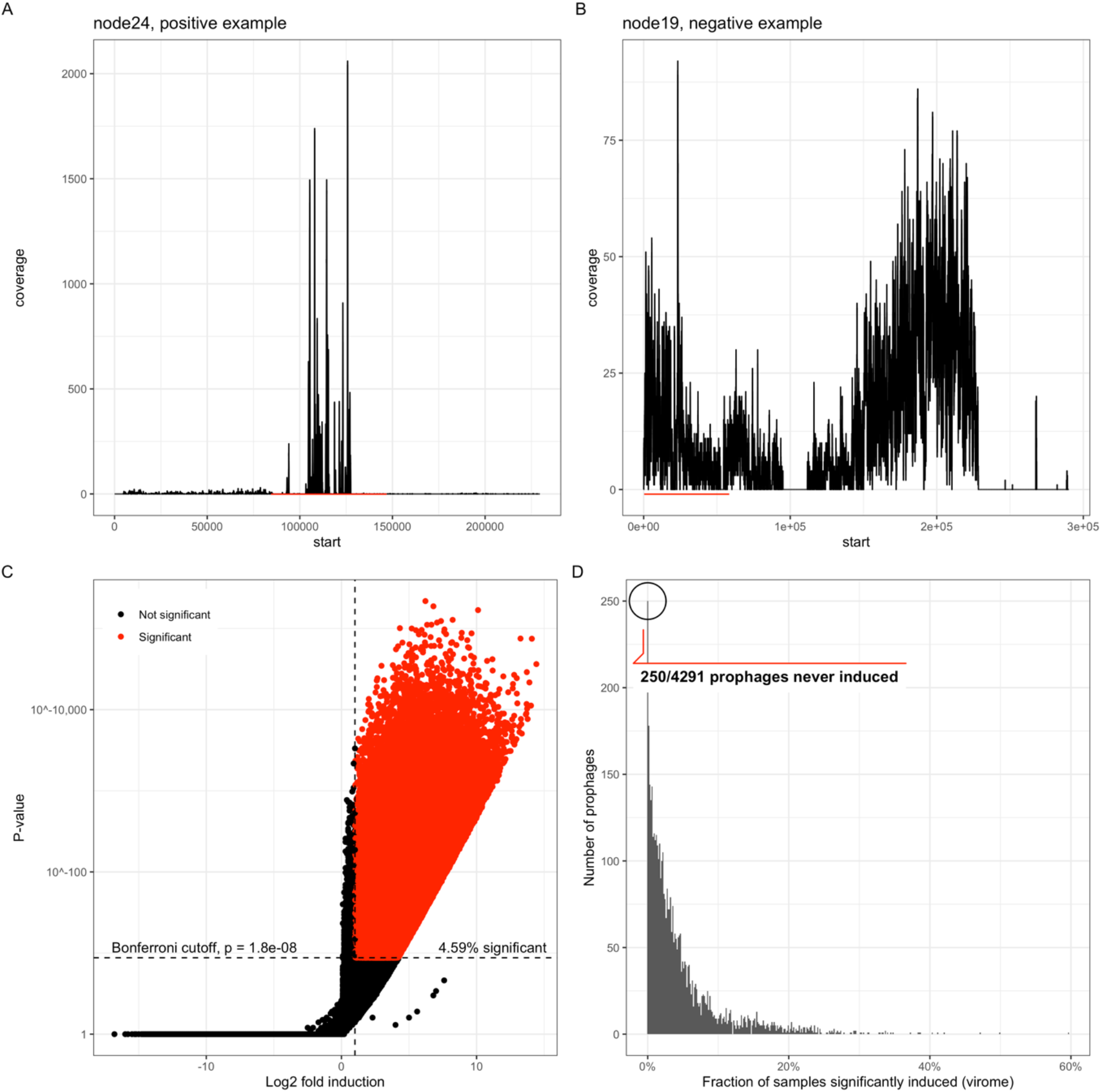
Evaluating induction in prophages using virome read mapping. A+B) Looking across the entire prophage carrying contig, we assessed induction as differential virome read coverage inside the predicted prophage region (red lines) versus the rest of the contig, which was considered as background. These two examples were chosen to illustrate this phenomenon as a positive (A) and negative (B) example. C) Volcano plot showing log2-fold induction vs. p-value (double log scale) distribution of all prophage/sample pairs. All prophage/sample pairs with an induction value > 1 and passing the Bonferroni cutoff were considered significant (red dots), which comprised 4.59% of the entire set. The red area looks larger due to massive overplotting in the lower part of the panel. D) Histogram of all prophages by how many children they were significantly induced in, ranging from 0% (no children, 250 prophages) to ~60% of all the children.

Comparing prophages across different predicted hosts, we found differences in the fraction of induced samples (**Fig 5A**, log linear model p < 2e-16). Among the genera with highest rates of induction were *Blautia, Bifidobacterium*, and *Erysipelatoclostridium*. Similarly, we found similar patterns of induction across prophages belonging to the same viral cluster (**Fig 5B**), in terms of which samples they were significantly induced in, suggesting that specific clusters of prophages are induced together within a sample.

**Figure 5:**
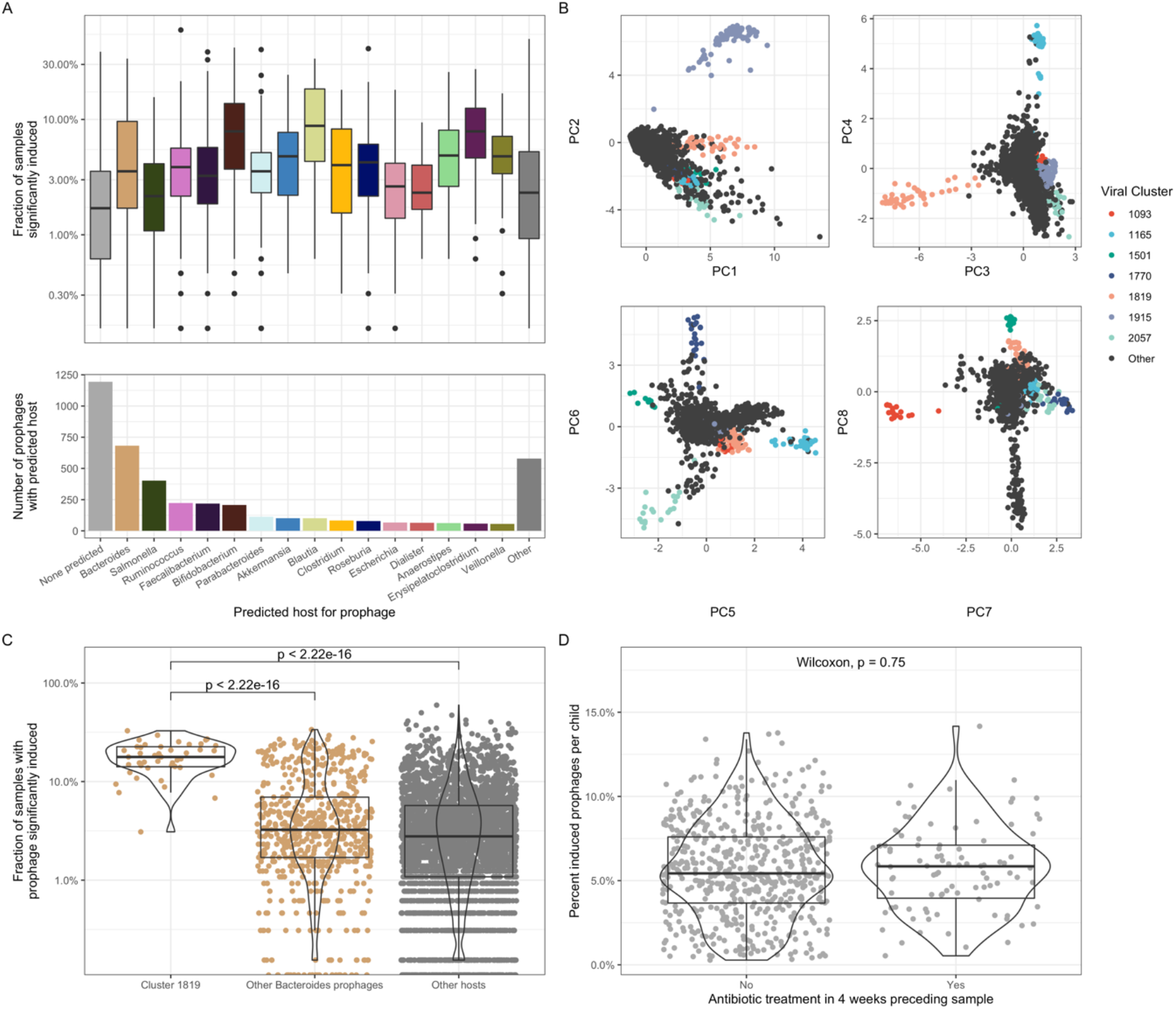
Comparing induction frequencies across vOTU and sample characteristics. A) Induction fraction by predicted host of the prophages, showing sizable differences between hosts (overall p < 2e-16), even when adjusted for mean RPK of the prophages. B) Principal Component Analysis (PCA), each dot is a prophage ordinated according to their induction pattern, ie. which children these prophages were induced in. Points which are close together signify prophages that tend to be induced in the same children. Points were colored according to some of the major Viral Clusters that show homogenous inductions patterns, notably including cluster 1819. C) Highlighting the induction frequencies of cluster 1819 prophages are much higher than other prophages with Bacteroides as their predicted host, as well as all other prophages with different predicted hosts. D) Children did not differ due to recent antibiotic exposure in terms of how many prophages that were found to be induced in their samples.

The most abundant group of prophages in the metagenomes, cluster1819, was among the most commonly induced (**Fig 5C**). Comparing the cluster against the rest of the predicted *Bacteroides* phages showed that the phages belonging to cluster1819 were more often significantly induced than the rest of the group, and to all prophages with different predicted hosts.

The significance of induction was also associated with the mean RPK of the prophages (Supplemental fig 1), however adjusting for this did not change the conclusions above. The use of antibiotics in the 4 weeks prior to sampling did not result in any significant difference in the proportion of induced prophages per child (**Fig 5D**).

## Discussion

This work has sought to characterise prophages in the infant gut in greater detail than has previously been achieved, and to highlight novel aspects of the associated phage biology. By combining machine learning based approaches we maximised our ability to identify integrated prophages from metagenomic assemblies. Of those identified, no single prophage could be found in more than 70 % of the samples suggesting that there is no ‘core virome’ of prophages in the infant gut. Much debate has taken place previously over the existence of a ‘core virome’ with early work on the topic identifying 23 phage contigs shared by at least 50% of samples from 62 individuals [74]; however, more recent studies with a larger sample size found that the most ubiquitous viral population was only present in 39% of the metagenomes used, and most of the populations were only sporadically detected at all [75]. This is something we see in our work corroborating their finding that most viral sequences were only sporadically identified throughout the samples; together these results suggest that infant viral communities are more unique to the individual than they are commonly shared. The infant microbiome has been shown to be a highly dynamic environment that constantly evolves until a more stable adult-like composition is reached around preschool age [1–5]. Considering this high degree of turnover, it may not be surprising to find prophages distributed sporadically throughout the samples as they adapt to changing bacterial host abundances, possibly reflecting ‘kill’ or ‘piggyback-the-winner’ dynamics [76, 77].

We were able to assign predicted hosts to ~65% of the prophages identified here, and these results found *Bacteroides, Salmonella*, and *Bifidobacterium* to be the most common hosts. *Bacteroides* and *Bifidobacterium* are well established as key members of the infant gut microbiota at one year of age in our samples [14] as well as in others [3, 4, 78], and so it would make sense for a large proportion of the prophages to infect these bacteria. The high proportion of *Salmonella* predictions may be an overestimation due to the tool used, which due to the high number of *Salmonella* spacers available may struggle to distinguish between *Escherichia* and *Salmonella* as predicted hosts and lead to an overprediction in the amount of *Salmonella*-infecting phages [63]. It is therefore important to remember that these are only predictions and whilst they largely make sense in the given environment there is no experimental evidence for hosts of these novel phages.

We also specifically looked at the prevalence of crAssphages, a well-known and large family of phages that are widespread in gut viromes [62]. In addition to the reference crAss-like phages that were used we also identified 109 additional vOTUs that clustered together and so were considered part of the crAss-like group. Our results strongly support previous work that has suggested that crAssphages are not abundant or prevalent in infants, and that this increases with age; they also support the suggestion that some of the crAss-like phages might be temperate [79].

Classification of the putative phages combined with the sequences of all sequenced bacteriophage genomes reported in the creation of 2934 clusters with 364 singletons. Of all the clusters identified, 953 were comprised solely of vOTUs identified in this work and of these 953 clusters, 177 were made up of a single member.

A more in-depth analysis of the most abundant cluster of prophages revealed an interesting perspective into their potential role in the infant gut setting. The most abundant group, Cluster1819, was comprised of 80 prophages closely related (genus or sub-family level) to *Bacteroides* phage Hanky p00’, a phage that was originally identified as an integrated prophage of *Bacteroides dorei* from metagenomic data [69]. Hankyphage was predicted to be present in half of the human population from geographically distant regions, and was found to lysogenize at least 13 different species of *Bacteroides* [69]. Our work utilises a large infant cohort study to further support the view that this group of phages is not only present in a large proportion of the population, but they are also abundant in these individuals.

The broad host range of these Hankyphage and other isolates in the work by Benler *et al* was found to be due to the possession of diversity generating retroelements (DGRs) that target tail fibres [69]; this was also something we identified in this work. Over half of the members of the viral cluster identified here were found to contain at least one DGR, all of which were predicted to target tail fibres. An increased diversity generated in the tail fibre sequence could lead to a host expansion as found originally, and therefore suggests that the most abundant group of prophages in the infant gut may have the potential to infect different strains of predicted host *Bacteroides*, a tactic that may be vital for their success in this dynamic environment. The relatives of Hankyphage identified here were also predicted to infect a few different hosts; whilst this may be an artifact of the host prediction method used it may also reflect an expansion of host range due to changing tail fibres. Without experimental evidence this cannot be validated but is an interesting possibility.

In addition to harbouring DGRs, members of the cluster also possessed a number of morons that may prove beneficial to their host or influence bacterial metabolism in some way. In this cluster we found evidence of genes involved in the dTDP-L-rhamnose biosynthesis pathway, which is responsible for biosynthesis of the O antigen of lipopolysaccharide in Gram negative bacteria [80–82]. *Bacteroides* in particular are known to produce a number of phase-variable capsular polysaccharides (CPS) [83, 84]. It has recently been shown that the expression of these CPS plays a role in the phage-host interaction of *Bacteroides*, being responsible for host-tropism of targeting phages [83]. The phase-variable expression of these CPS creates diversity within the otherwise homogenous population of *Bacteroides* and this may help to ensure their survival by maintaining subpopulations that are differentially susceptible to phage infection, and also making them better equipped to survive environmental changes [83]. The possession of genes involved in the LPS biosynthesis pathway is an established mechanism of some phages such as *Pseudomonas aeruginosa* phage D3 and *Streptococcus thermophilus* phage TP-J34 to ensure superinfection exclusion of other phages [33, 34]. In this way we probably witness a piggy-back-the-winner model of infection, something that has recently been proposed for the crAss-like phage crAss01 and its *Bacteroides* host [85].

This is, however, only one of a number of reasons as to why these phages could possess genes in this pathway. In addition to being a receptor site for the attachment of phages the O-antigen is of importance for recognition of the human immune system and the pathogenicity of the bacterium [86, 87]. Modification of the O-antigen in some instances has also been shown to affect the bacteria’s ability to establish infection through its ability to bind to epithelial cells, such is the case in *Pseudomonas aeruginosa* containing the FIZ15 prophage [88].

Another gene of interest found in the cluster was *menA*: a component of the menaquinone biosynthesis pathway. Menaquinone (vitamin K2) is an important part of the electron transfer pathway in prokaryotes and is vital in humans in the blood clotting process, and bone and nervous system health [89–93]. Whilst the importance of microbially synthesised vitamin K is debated due to the low amount of total vitamin K it would be contributing [94–96]. This may be different in infants where there are risks of deficiency in early life due to the limited diet [97]. In these instances, the microbially synthesised portion may be of more importance: *Bacteroides* in particular are known to be one member of the gut microbiota that produces menaquinone [89], and so the presence of a component of the pathway in a *Bacteroides* prophage suggests that they could also influence production.

Our induction analysis shows that out of the subset of phages we were able to test, most prophages that were identified as present in this work were also induced at least once. This suggests that these phages are an active part of the community and may play a prominent role in the shaping of the bacterial community in the gut. Previous work has suggested that the infant gut in particular may be dominated by temperate phages that may be induced due to the high turnover rate/constant maturation of the bacterial community during the first few years of life [19, 20, 22].

Environmental conditions can lead to the induction of prophages from their bacterial hosts, leading to lytic replication and the production of progeny phages. A number of factors have been shown to induce prophages such as certain chemicals (mitomycin c) and antibiotics such as fluoroquinolones [98–100]. More recently, the use of common oral medications such as nonsteroidal anti-inflammatory drug diclofenac, and other antibiotics including ampicillin, norfloxacin, and ciprofloxacin were shown to induce prophages from bacterial isolates of the human gut [101]. Whilst our work showed no effect of antibiotics on the proportion of induced prophages this may be due to time limitations of the method. Evidence of induction may have already been turned over in the 4 weeks preceding sampling, and this may be why we cannot see it here. The free phages produced by induction may have been broken down in the environment in this amount of time. This highlights the need to be careful of interpreting the ‘snapshot in time’ of a population from a single time point and indicates the need for more longitudinal data for work such as this.

## Conclusions

In summary, our results show that prophages of the infant gut form a diverse community that is different in each individual; no conserved ‘core’ virome of temperate phages was apparent. Our work utilises a large infant cohort to support the previous observation that crAss-like phages are present in small numbers early in life and become more dominant later. We also identified a novel cluster of phages that are the most abundant in the metagenomes, which are genetically similar to *Bacteroides* phage Hanky p00’. The possession of DGRs targeting tail fibres in members of this cluster suggest they may be able to infect a range of bacterial hosts. We also found evidence that they may modify host LPS through possession of components of the dDTP-L-Rhamnose pathway. Therefore, this group of phages possess elements that may allow them to maintain differentially susceptible subpopulations of their host bacterium, whilst also containing DGRs that could expand their host range. By utilising the paired metagenome and virome sequencing we were able to show that out of those phages we were able to test, the majority of them were induced at least once. However, testing induction against antibiotic usage revealed no association between the two factors, although this may be a reflection of the speed at which the evidence of induction is turned over in the gut. This highlights the need for more longitudinal data in the field to ensure that associations are not based on a single moment in time, that may not be representative.

