## Supplementary Figure 1 for "Prophages in the infant gut are largely induced, and may be functionally relevant to their hosts"

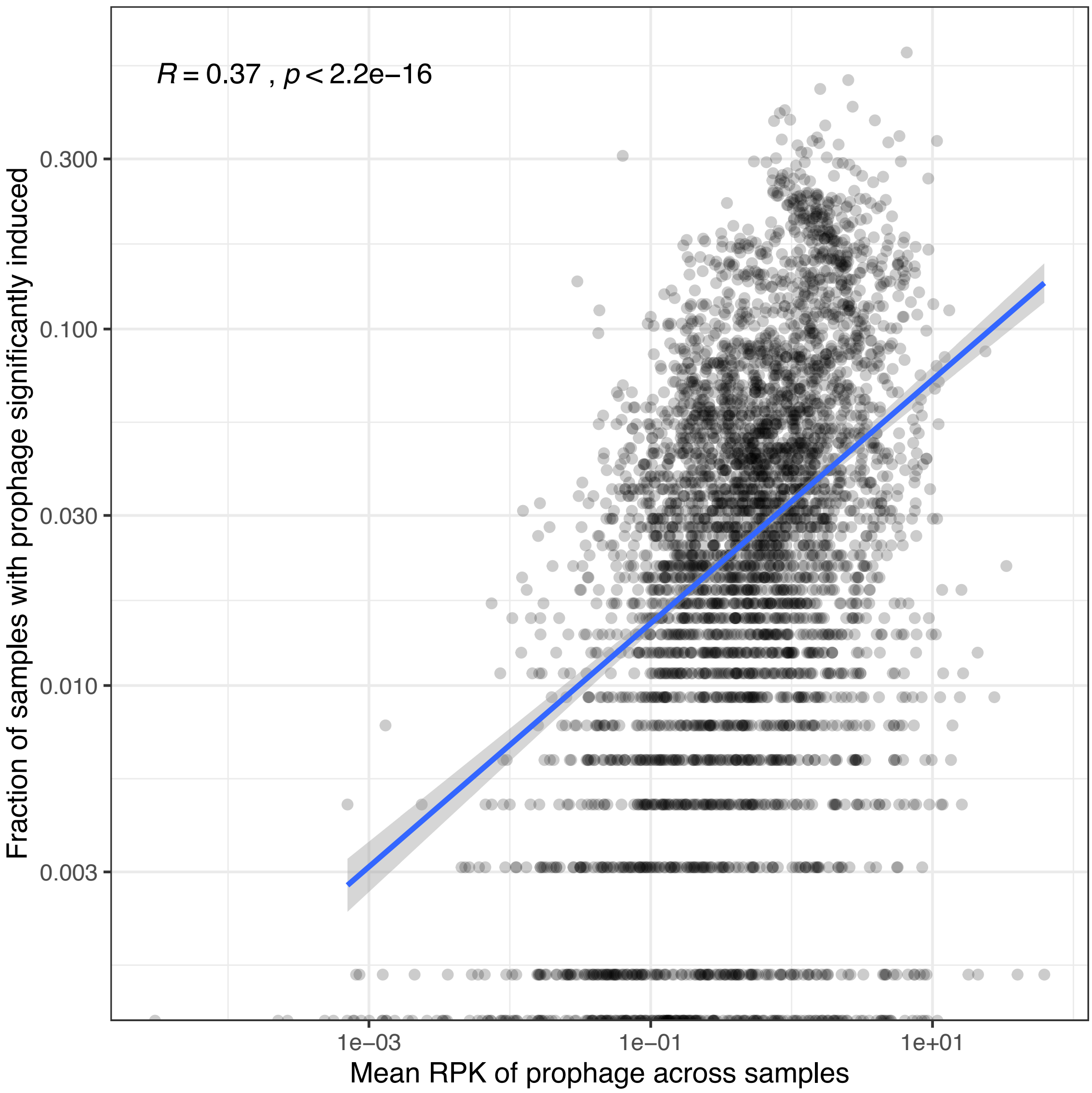


***Supplementary Figure 1:*** *There was a strong correlation between the abundance of a prophage (x-axis, measured as mean Reads Per Kilobase (RPK) across all samples, log scale) and the proportion of samples with significant induction (y-axis, log scale), quantified with Spearman correlation of 0.37, $p < 2.2e-16$. Therefore, all analyses of proportions of significantly induced samples per prophage were adjusted for the mean RPK.*
